# The chemistry of families: Linking body odor composition to olfactory perception

**DOI:** 10.64898/2026.09.29.755001

**Authors:** Katharina Hierl, Carolin Schnauber, Marlen Kücklich, Bastian Heinlein, Claudia Birkemeyer, Brigitte M. Weiß, Anja Widdig, Laura Schäfer

## Abstract

Body odors inform human kin recognition. However, underlying chemical mechanisms and how they relate to body odor perception are still unclear. The present study therefore simultaneously examined perceptual body odor similarity according to human raters, its chemical basis and the link between the two in human families. The study included 33 raters who assessed body odors collected from 60 odor donors belonging to 15 biological families. Each family comprised a mother, a father and one adolescent same-sex sibling-dyad (8 sister dyads, 7 brother dyads). Body odor sampling was conducted with cotton pads for presentation to human raters and with thermal desorption (TD) tubes for Gas Chromatography–Mass Spectrometry (GC-MS) analysis. Relatedness and sex predicted perceived body odor similarity in parent-offspring versus unrelated control dyads. Analyses of chemical profiles revealed greater body odor similarity within families than between unrelated individuals. Parallel to the perceptual results, relatedness and sex predicted chemical similarity based on whole odor profiles. Finally, a machine-learning prediction model integrating perceptual and chemical data, revealed a small but significant positive association between chemical similarity of body odor and perceived olfactory similarity according to human raters, suggesting that odor perception is partly informed by chemical characteristics. Overall, the present study extends our understanding of human chemosensory communication and supports the notion of a family-specific olfactory signature.

## Introduction

According to the *Kin Selection Theory* (1), prosocial behavior develops as a function of evolutionary benefit and is expressed towards close relatives. Kin recognition is associated with kin biased behavior, for example, selective parental investment (2,3), preferred social relationships (4) and incest avoidance (5) in humans and non-human primates. Several mechanisms of kin recognition have been proposed. Phenotype matching is one prominent mechanism (reviewed in 6) and is characterized by deriving relatedness information from phenotypic traits, such as visual, acoustic and olfactory cues (7,8). Individual body odor is one such major feature recognized in humans and non-human primates (9,10).

Studies on non-human primates indicate that kinship information is reflected in chemical profiles: In rhesus macaques (*Macaca mulatta*), related animals exhibit higher similarities in their chemical profiles, especially close maternal relatives (i.e., maternal half-sisters (11). Similar findings were observed in catta lemurs (*Lemur catta*; 24) and chimpanzees (*Pan troglodytes*; 9).

In humans, third parties are able to match mother-infant dyads (13) and homozygous twins (14) on the basis of their body odor alone. These findings suggest the existence of a family-specific body odor signature that may serve as an olfactory phenotype detectable by unrelated individuals. Since body odor is composed of a complex mixture of hundreds of volatile organic components, assessment of family-specific clustering of these components is a promising approach to capture the basis of olfactory kin recognition. Yet, to date, little is known about whether the chemical body odor profiles of biologically related individuals are more similar to one another than to those of unrelated individuals, or to what extent similarity in body odor might be expected within biological families.

Data on chemical body odor similarity between human relatives are so far limited to homozygous twins: Kuhn & Natsch (14) observed that odor profiles of twin pairs overlap (51.5% variance explained by sibling pair), confirming that body odor is strongly determined by genotype (15). It thus can be assumed that other dyads of the nuclear family, such as parent-child-dyads, also share similar body odor profiles. However, no data are currently available on this. Further, the question remains whether and how the chemical composition of a family-specific olfactory signature corresponds to the olfactory percept.

The present study hence aims to fill these gaps by systematically examining the interplay between body odor perception and chemistry in human biological families: We therefore target the following research questions: In the perception study, we investigate (i) the perceived body odor similarity of related dyads (parent-child dyads, full-sibling dyads) assessed by third-party raters, whereby we expect greater perceived body odor similarity for related pairs and for same-sex pairs (H_p_1). In the chemical study, we investigate ii) the chemical similarity in body odor of related dyads (parent-child dyads, full-sibling dyads) by GC-MS analysis, expecting larger family-wise body odor profile similarity among biological families compared to unrelated individuals (H_c_1), dyad-level greater chemical similarity in related vs unrelated dyads as well as in same-sex dyads (H_c_2). Finally, we explore iii) the relationship between perceived and chemical body odor similarity by adopting an integrative neural-network model, expecting a positive association between the two (H_pc_1).

## General Methods

### Ethics

The ethics committee of the University of Dresden (Code: SR-EK-477112022) approved this study in accordance with the ‘World Medical Association’s Declaration of Helsinki’. Written, informed consent was received from all participants. The study was pre-registered at DRKS (German Clinical Trials Register [Deutsches Register Klinischer Studien] trial number: DRKS00032290, see *Figure 1* for an overview of the study).

**Fig 1.**
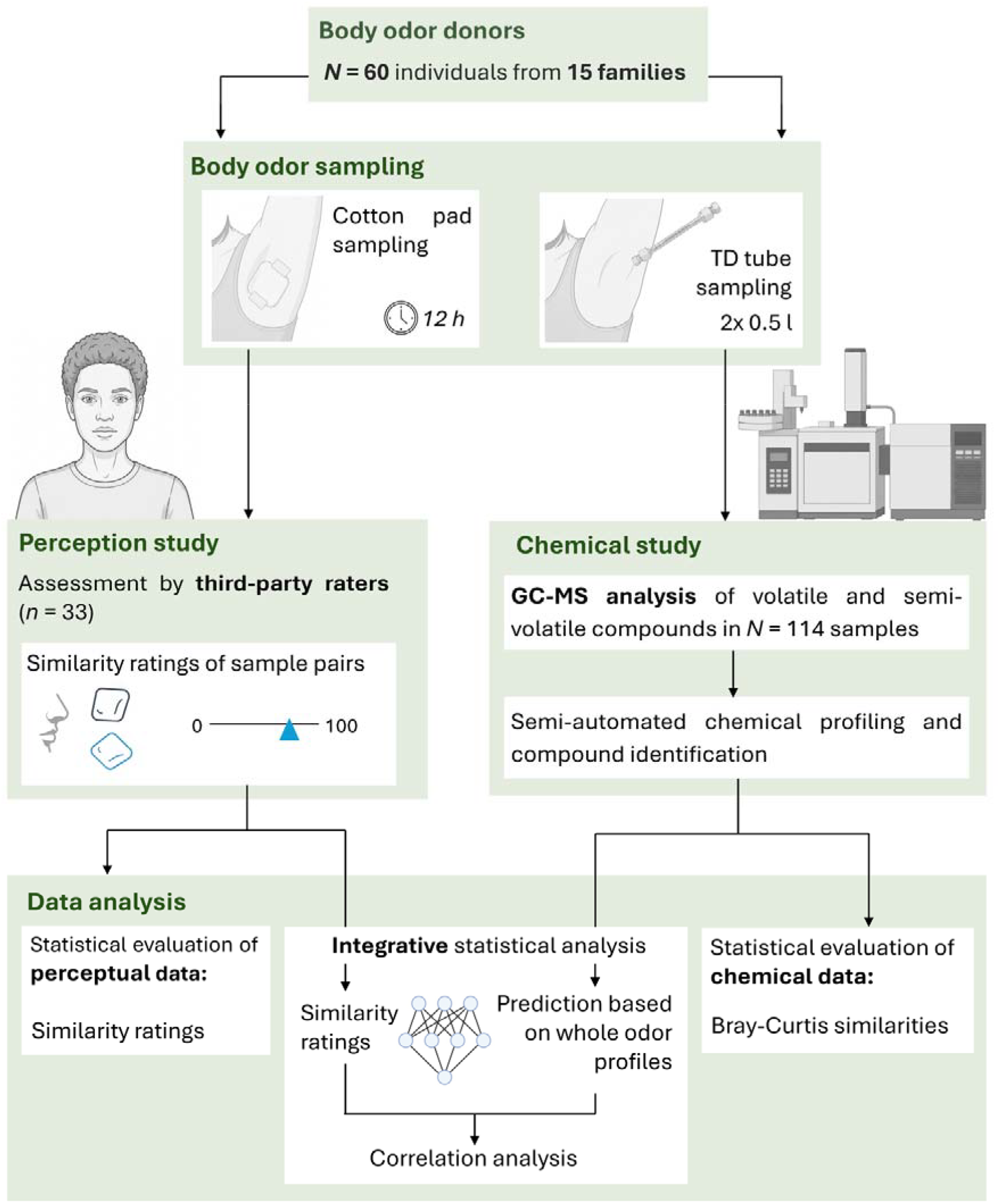
Overview of the study procedure, including sample and data collection.

### General sampling approach

The body odor donor sample comprised *N* = 60 participants from 15 families with 4 members each. Each family included a) mother (*M*_*age*_ = 46.20; *SD* = 2.83 years), b) father (*M*_*age*_ = 48.85, *SD* = 4.18 years), c) child 1 and d) same-sex child 2, resulting in a total of 8 families with daughter dyads (*M*_*age*_ = 15.94, *SD* = 1.65 years) and 7 families with son dyads (*M*_*age*_ = 15.93, *SD* = 2.46 years). Inclusion criteria were biological relatedness of family members (according to self-report by parents), normosmia, and sharing of one household. To control for developmental fluctuations of body odor related to kin recognition (10,16), children had to be between 12 and 20 years old and in late- or post-pubertal developmental stage (assessed by the Pubertal Development scale, PDS (17); daughters: *M*_*PDS*_ = 11.44, *SD* = .81; sons: *M*_*PDS*_ = 9.21, *SD* = 1.76). Meanwhile, exclusion criteria were regular drug use, and severe chronic or hormonal diseases.

At an initial appointment, body odor donor families were equipped with a study kit containing all materials required to collect cotton-pad body odor samples, and respective instructions. Participants were asked to adhere to a behavioral protocol 48 hours before both sampling with cotton pads and thermal desorption (TD) tubes at T1 and T2 (*see chemical study*). This involved avoiding odor-intensive foods, alcohol, following a vegetarian or vegan diet (18,19), and no use of perfumed products for clothing and bedding (see supplement, *body odor sampling procedure*). The protocols for sampling with cotton pads, and TD tubes are described in the perception study – and chemical study section, respectively.

### Perception study

#### a) Body odor sampling procedure

##### Body odor sampling with cotton pads for assessment by third-party raters

To obtain body odor samples with cotton pads, participants were asked to adhere to a standardized behavioral protocol that has been established in numerous studies (16,19,35; see supplement, *body odor sampling procedure)*. Cotton pads were attached with medical adhesive tape in the evening and worn under both armpits for 12 hours throughout the night. At the end of the sampling period, cotton pads were placed back in air-tight pharmacy jars (30 ml, wide-mouth) and additionally sealed with PTFE isolating tape. Within 10 h, body odor samples were frozen at −25 °C at the University Hospital Dresden and stored until they were needed for perception experiments (22).

#### b) Body odor raters

Third-party raters were *N* = 33 adults (16 females, 17 males) with normal olfactory ability, aged 22 to 35 years (*M*_*age*_ = 28.09, *SD* = 3.46). Olfactory ability of all human raters was tested using a screening version of the Sniffin’ Sticks (37; supplement, *rating procedure*). In addition, current olfactory impairments (e.g. rhinitis) were queried at the beginning of the appointment to ensure intact olfactory functioning, and assessment was postponed if impairments were present.

#### c) Rating procedure

In order to reduce contamination with other smells, no perfumed products or any food items were allowed to be used in the laboratory in which all ratings took place. The laboratory was aired between individual sessions. Furthermore, the experimenter refrained from wearing perfumed products and smoking and wore nitrile gloves. Likewise, third-party raters were asked to refrain from wearing perfume and other strong-smelling care products, from smoking on the day of the experiment, and from eating and drinking coffee 1 hour before the experiment. Third-party raters were invited to evaluate body odor samples *(see Fig 1*, body odor samples (“How similar do these two body odor samples smell?”) on a visual analogue scale from 0 to 100 (0 = not similar at all, 100 = completely similar). Related and unrelated dyad-combinations from a total of 3 families (*n* = 12 individuals) matched for sex of the children were presented to each third-party rater. The experimental order was randomized, and a total of 20 pairs were assessed by each rater.

#### d) Statistical analysis

All behavioral data were analyzed using Python, version 3.9.22 (24).

##### 1) Perceived body odor similarity (H_p_1)

We analyzed similarity judgements using mixed–effects models with gaussian error distribution to account for the hierarchical data structure with repeated measurements nested within raters. Separate models were estimated for parent-child, and sibling dyads. Models were estimated using maximum likelihood as implemented in statsmodels (version 0.14.4). Inspection of residual diagnostics (histograms, normal Q– Q plots, and residuals-versus-fitted plots) and Q–Q plots of the random intercepts indicated deviations from the Gaussian assumptions. To obtain robust inference, we therefore additionally performed cluster bootstrap resampling at the participant level (1,000 replicates). Bootstrap estimates, percentile confidence intervals, and empirical two-sided p-values were computed for each fixed effect. There is no cause for concern regarding multicollinearity, as all Variance inflation factors (VIFs) were below 2. see supplement, *rating procedure*). They were asked to rate the perceived similarity of two

1. For parent-child and control adult-adolescent dyads, we specified a linear mixed model (LMM) with relatedness (0 = not related vs. 1 = biologically related), and opposite-sex vs. same-sex (0 vs. 1) as fixed effects. A random intercept for third party-rater ID was included to model within-subject effects.
2. For siblings and sex-matched control dyads, we fitted an LMM including related vs. not related (0 vs. 1) and sex of the dyad (female vs. male) as fixed effects, again with a random intercept for rater ID.

#### e) Results

##### 1) Perceived body odor similarity: effect of relatedness and sex (H_p_1)

A. For parent-child and respective control dyads, results indicated that perceived similarity was higher when dyads were related (β = 8.74, SE = 2.41, *z* = 3.63, *p* < .001, 95% CI=[4.05; 14.27]), and when they were of the same sex (β = 5.26, SE = 2.38, *z* = 2.21, *p* = .027, 95% CI = [1.05; 9.07]; *see Fig. 2*). Between-rater variance was modest (σ^2^ = 35.1) relative to within-rater variance, indicating limited but non-negligible clustering effects.
B. For sibling dyads and adolescent control dyads, relatedness and sex did not predict perceived similarity (relatedness: β = –5.22, SE = 4.68, *z* = –1.12, *p* = .265, 95% CI = [–14.40; 3.95]; sex: β = –2.659, SE = 5.24, *z* = –.51, *p* = .612, 95% CI = [–12.93; 7.61]).

**Fig 2.**
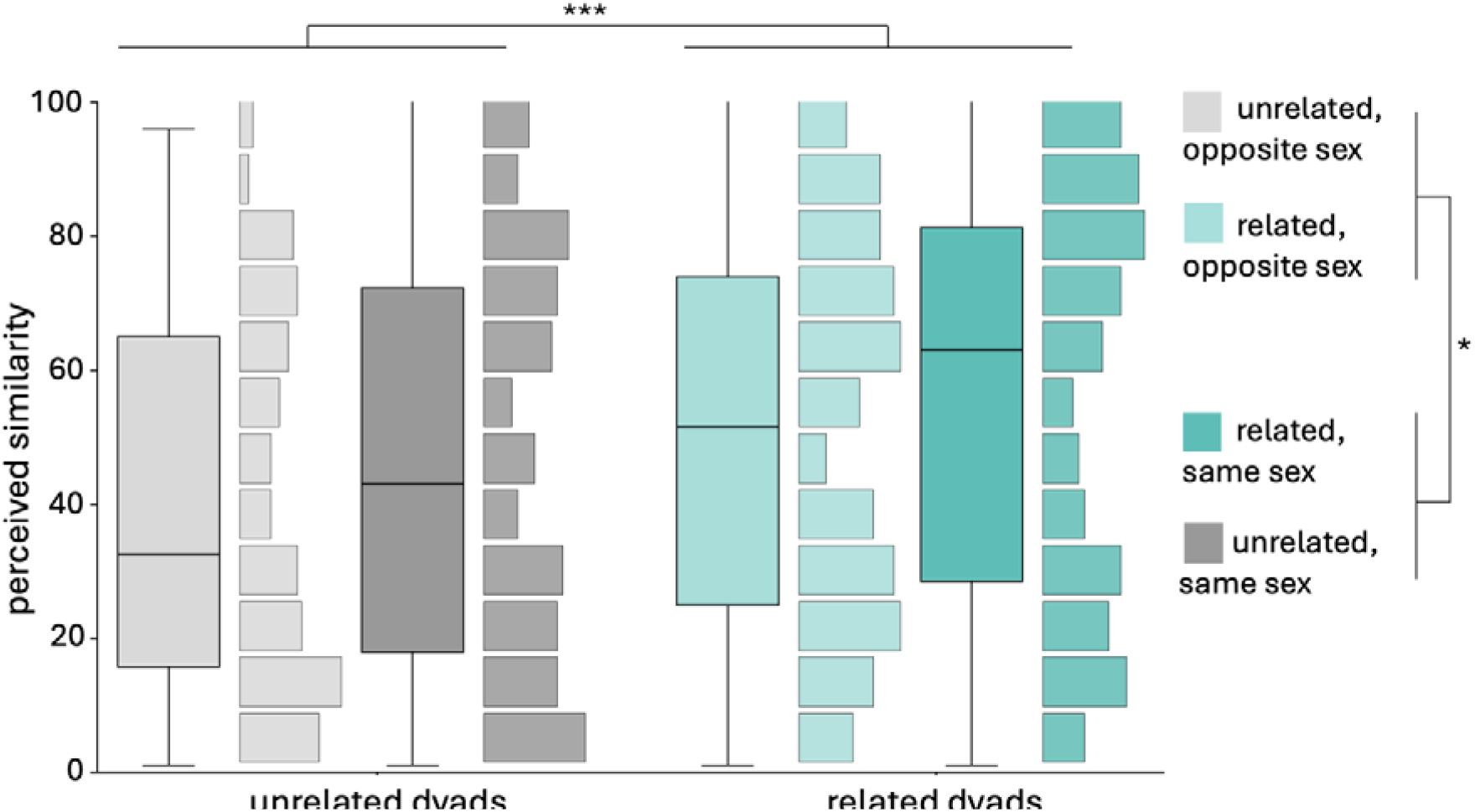
Perceived similarity of related parent-offspring dyads and unrelated control dyads according to human raters. * p <.05, *** p < .001.

### Chemical study

#### a) Body odor sampling procedure

##### TD tube sampling for chemical analyses

At the same appointment collecting samples for perception study in the lab (T1), body odor components were sampled for the chemical study using stainless steel TD tubes (Supelco 1/4 in. × 3 1/2 in., Supelco, Bellefont, USA). A second TD tube sampling point (T2) was conducted at least three days after (*M* = 19.60, *SD* = 18.62, min = 3, max = 61 days). TD tubes are adsorbent traps, which, in our study, contained two polymers (.095 g Tenax TA and .21 g XAD-2, Sigma Aldrich) to capture the volatile (VOC) and semi-volatile organic compounds (sVOC) emanating from the body. Combining these two polymers ensures comprehensive coverage and sensitivity for sampling VOC (substances that evaporate easily, pass into the air and can be directly detected by the main olfactory system) and sVOC (substances that are less volatile than VOCs and are more likely to be found directly on surfaces such as skin; 40,41). For sample collection, each TD tube was positioned as close as possible to the axillary of the participant without touching their skin. TD tubes were connected to an air pump (BiVOC2, Holbach) with a silicone hose. The air pump was set to a constant 1.5 L/min flow rate until .5 L of air was collected. For each participant, samples were taken from the left or right armpit in random order at T1 and T2, respectively. The experimenter wore a lab coat, medical mouth-nose cover and nitrile gloves to minimize contamination. In addition to the body odor samples, 11 room blanks (i.e. .5 L of air while only experimenter was present, and after airing the room for 30 min), and 3 analytical blanks (i.e. unopened TD tubes, otherwise handled like the other TD tube samples) were collected as control for contaminations. Both sampling methods are summarized in *Fig* 1 (and see supplement, *TD tube sampling*).

#### b) Chemical analyses

##### Gas Chromatography – Mass Spectrometry (GC-MS) analysis

Analysis of chemical body odor samples was conducted using a GCMS-TQ8040 consisting of a gas chromatograph GC-2010 Plus and a triple quadrupole mass spectrometer. A thermal desorption system TD-20 (Shimadzu, Kyoto, Japan) was used for sample introduction. Samples were injected at a split ratio of 1:5 and desorbed for 8 min at 250°C and a helium flow of 60 mL/min to a Tenax TA-filled cold trap (−20°C). For subsequent injection, the cold trap was heated up to 250°C under a helium flow of 14.5 mL/min to release the trapped compounds into the GC. Electron-impact ionization was accomplished at 70 eV and 200°C with a scan range of mass-to-charge ratio (*m/z*) of 30 - 500. Starting at 35°C for .5 minutes, the temperature increased at a rate of 6°C per minute until reaching 320°C. This temperature was held for 15 min which resulted in a total program time of 63 min. The final sample number comprised *N* = 114 excluding blanks (see supplement for details on measurements of samples).

##### Chemical profiling

The resulting chemical profiles were processed with a semi-automated method using AMDIS (v 2.65, 42) and Shimadzu GCMS Browser (see supplement, *chemical profiling*). After exclusion of substances that were abundant in room blanks, the final dataset consisted of 130 substances (an amount similar to other studies on non-human primates, see 43). For analyses, relative peak areas (peak area/(sum peak areas of included substances) were used.

#### c) Statistical analyses

All chemical data were analyzed with R, version 4.4.0 (29).

##### 1) Chemical body odor similarity *within* vs. *between* biological families (H_C_1)

A nonparametric Analysis of Similarity (ANOSIM) (package ‘vegan’, version 2.6-2; 45) based on pairwise Bray-Curtis (BC) indices (31) was conducted to determine the overall similarities between whole odor profiles (all 130 substances) of families, controlling for repeated measures by setting individual ID as stratum. Peak areas were standardized and log(x+1)-transformed for the analysis. The ANOSIM was performed with a customized R script strongly paralleling the one by Oksanen and colleagues (45), investigating similarity distances (BC similarities) within and between families while controlling for the Sample ID.

##### 2) In-depth assessment of chemical similarity: effect of relatedness and sex (H_C_2)

Similarities of whole body odor profiles were further investigated using multiple membership multilevel models (MMMM) which enable the handling of data structures with multiple affiliations. For instance, a sample ID of a daughter from “family_3” might occur in two different columns – one for the comparison of that daughter with her mother, and one for the comparison of that daughter with an unrelated adult woman, while affiliating these data points with the same ID. Furthermore, MMMM are suitable for the analysis of data that are not strictly hierarchical (32). These models were also based on pairwise Bray-Curtis (dis)similarities calculated using the package ‘vegan’, version 2.6-2 (45) from standardized, log(x+1)-transformed peak areas which were computed previously. The continuous predictors age and sampling date were z-transformed. Given that similarity scores follow a beta distribution, all models were calculated using a Bayesian approach in the package ‘brms’ (version 2.21.0, 47), a package able to handle multi-membership data with beta distribution, with resulting estimates referring to the posterior distribution. Bayesian estimation was used exclusively for the multiple-membership beta regression models because this modeling framework is currently most readily available in a Bayesian implementation, whereas all remaining analyses were conducted using frequentist methods.

We first examined effects of relatedness on BC-similarity in i) parent-child versus unrelated control dyads, and ii) siblings versus unrelated dyads with family ID as test predictor. Next, effects of sex on BC-similarity were examined in iii) same-sex parent-child versus opposite-sex parent-child dyads with sex (same) as test predictor, and iv) sister-dyads versus brother-dyads with sex (female) as test predictor (see supplement for details on model specifications).

#### d) Results

##### 1) Chemical body odor similarity *within* vs. *between* biological families (H_C_1)

Comparing the chemical body odor profiles of 15 families revealed that profiles of individuals from the same family were more similar than profiles of individuals from different families (ANOSIM, N=114, *r* = .1927, *p* = .001; *Fig. 4a* depicts all included samples).

##### 2) In-depth assessment of chemical body odor similarity: effect of relatedness and sex (H_c_2)

MMMM revealed an effect of relatedness when comparing chemical similarity scores of parent-child vs. unrelated dyads indicating higher body odor similarity in related parent-child dyads (coef. = .213; *SE* = .046; CI = [.123; .303]; see *Fig. 4b*). Similarly, siblings exhibited greater chemical similarity: when compared to unrelated dyads, the MMMM indicated an effect of relatedness (coef. = .211; *SE* = .07; CI = [.075; .349]; see *Fig. 4b*). MMMM comparing same-sex versus opposite-sex parent-child dyads also indicated an effect of sex showing greater chemical similarity in same-sex dyads (coef. = .156; *SE* = .08; CI = [.001; .312]). Comparing similarity scores between brothers and sisters, the MMMM indicated that sex of the sibling dyad had no effect on chemical similarity (coef. = .006; *SE* = .147; CI = [-.281; .302]).

**Fig. 4.**
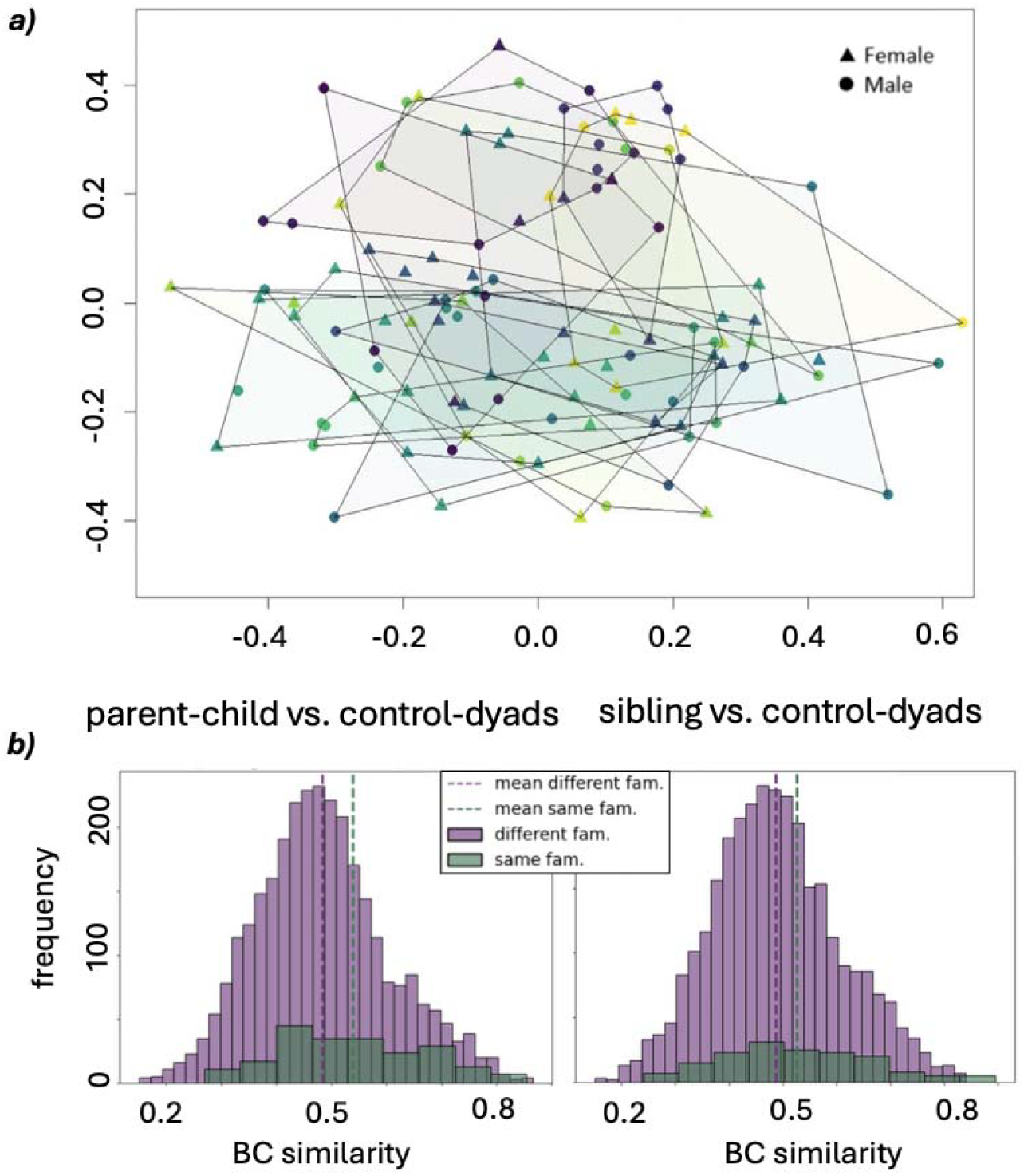
a) Two-dimensional non-metric multidimensional scaling (NMDS) plot of chemical body odor profiles of human families. Each icon represents one individual. The axes have arbitrary scales and closely positioned data points represent similar chemical profiles. Color symbols connected via a line depict family affiliation. b) Distribution of BC-similarity with respect to the expected relatedness in parent-child vs. unrelated dyads and sibling vs. unrelated dyads.

### Integrative analyses

#### a) Statistical analyses

##### 1) Relationship between perceptual and chemical body odor similarity (H_pc_1)

Finally, we investigated whether the similarity of two body odor samples as perceived by the third-party raters could be predicted from the chemical composition of those samples. Specifically, we hypothesized that the chemical profile data obtained from our GC-MS analysis predicted the perceptual similarity reported by the third-party raters (H_pc_1). To test this, we trained a machine learning model to predict the perceptual similarity between pairs of samples from the chemical profile data (see Figure 5 a). If the model could predict perceived similarity significantly better than chance, this would indicate that patterns in the chemical data are informative of olfactory perception. Recent studies revealed that the perceptual similarity between odorants or odorant mixtures with few components can be predicted using machine learning models (34,35). However, this has not yet been applied to complex odorant mixtures with more than 100 identified components, as in the present study.

**Figure 5.**
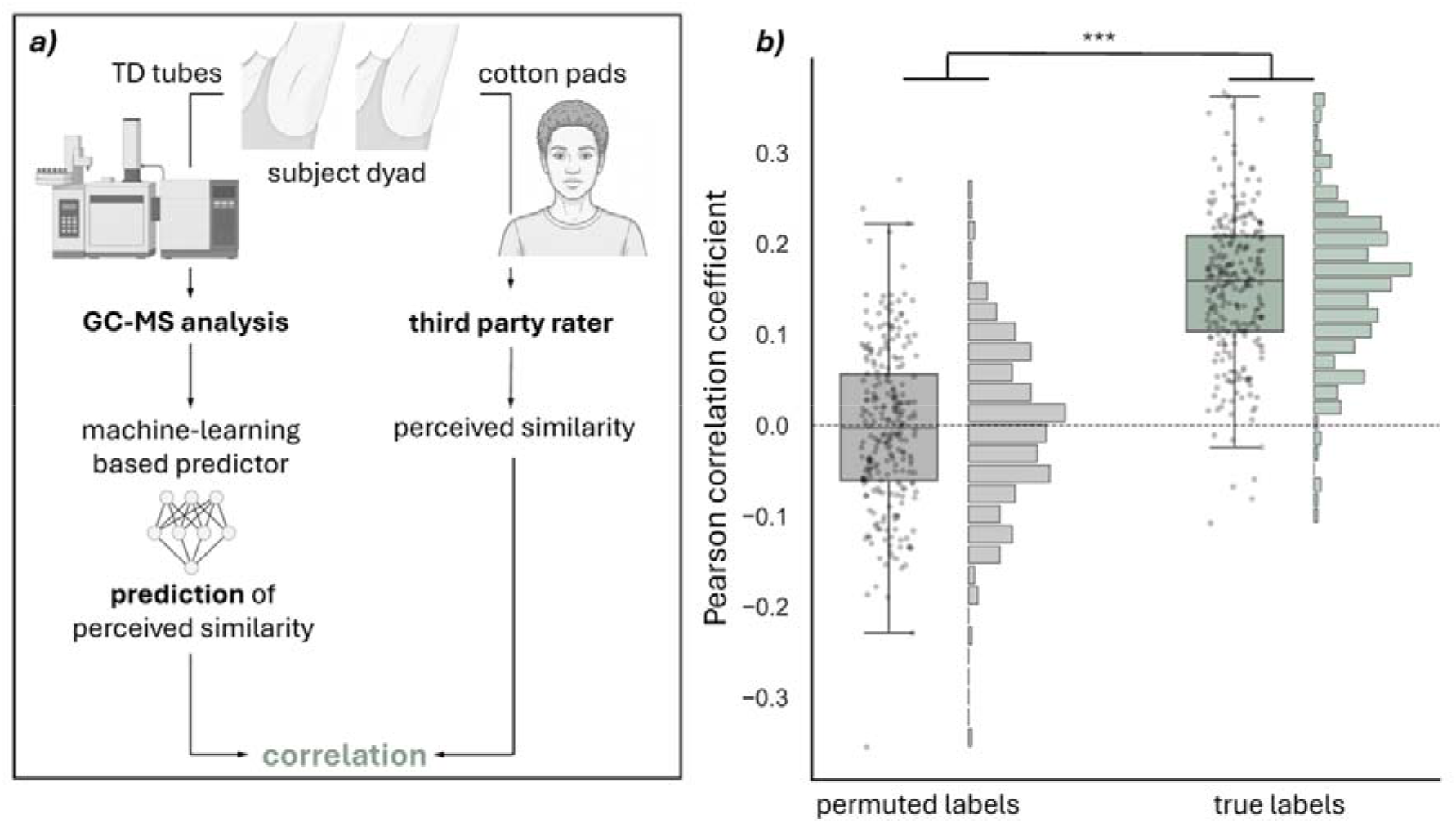
a) Approach of integrative statistical analysis. b) Correlation of predicted similarity ratings and similarity rating from third-party olfactory assessment. Permuted labels = baseline condition. True labels = target prediction. *** p < .001.

For our analysis, we built a dataset of 640 chemical-perceptual similarity sample pairs. Hence, each element in the dataset comprised the absolute difference vector of the log(x+ 1)-transformed peak areas obtained from the GC-MS analysis, and a perceived similarity rating. We normalized each difference vector to have unit-variance. Additionally, we normalized the ratings of each rater to have zero-mean and unit-variance. In accordance with standard practice in machine learning, we then split the dataset into training- and test set with 512 and 128 samples, respectively, to account for possible over-fitting on the training data (36). To evaluate the quality of the predictions of the machine learning model, we computed the Pearson correlation coefficient between predicted similarity rating and the true similarity rating in the test set. We repeated this *n* = 250 times with different sample assignments to training and test sets to avoid distortion of the correlation coefficient by a specific dataset split. To test whether perceived similarity could be predicted above chance, we compared the resulting distribution of correlation coefficients using a one-sided Mann-Whitney-U test to the distribution of correlation coefficients obtained when trying to predict randomly permuted similarity ratings in the test set. For further robustness, we additionally implemented a simple linear regressor and an individual-based dataset splitting procedure for prediction of similarity ratings. Details see supplement (ablation study).

#### b) Results

##### 1) Relationship between perceptual and chemical similarity (H_pc_1)

Averaged across all dataset splits (*n* = 250), model-predicted similarity was significantly correlated with perceived olfactory similarity (*r* = .152). In contrast, permuted similarity ratings were uncorrelated with perceived olfactory similarity (*r* = −.003). A Mann-Whitney-U-based comparison of the two correlation coefficient distributions revealed a significant above-chance predictive capability of the machine learning model (*p* < .001; Figure 5. b).

The additional analyses outlined in the supplement support this finding, as significant Mann-Whitney-U tests indicated that learning occurred in all scenarios. Thus, the integrative analyses imply a relationship between chemical and human perception data.

## Discussion

The present study integrated perceptual and chemical data to explore a family-specific olfactory signature and found evidence for human olfactory kin recognition based on chemical profiles, human perception and the computational integration of the two.

In the perception study, we observed higher perceived body odor similarity ratings for parent-child (vs. unrelated control) dyads, and for same-sex (vs. opposite-sex) dyads. According to our hypotheses (H_p_1), these findings confirm a signature of genotype and sex (15) to be reflected in similarity judgements. However, there was no such effect among adolescent siblings. This should not be interpreted as evidence for the absence of a familial chemical signature among siblings. Importantly, our family design provided substantially fewer unique sibling dyads compared to parent-child dyads: within each four-member family, only one sibling dyad was available, compared with four parent child dyads. Consequently, the sibling analysis may have provided less information to detect a potentially weaker similarity effect. Furthermore, whereas parent-child dyads always share 50% of the genome, the overlap in genome-wide similarity between human siblings can vary between 36 and 63% (37,38). Recent studies showed that higher genome-wide similarity increases kin bias and kin recognition in nonhuman primates (Freudiger et al. in prep), however, this was not controlled for in our sample. Future studies may therefore consider the actual genome to account for more fine-grained genetic effects on body odor.

Our results from the chemical study extend previous findings on homozygous twins (14) to full-siblings and parent-child dyads: Aligning with our hypotheses, chemical body odor similarity analysis provided evidence of a clustered representation of family-specific body odor, paralleling previous studies in non-human primates (H_c_1; 17,46). In-depth analyses further revealed that relatedness and sex predicted greater odor profile similarity (H_c_2). However, our present effect size is rather small (*r* = .19), indicating that body odor composition is not exclusively structured by relatedness, but also show overlaps across families. Hence, additional factors may attenuate the availability of reliable kin cues that may serve as the basis for body odor matching to promote successful olfactory kin recognition. The individual microbiome may be one of these additional sources of variance between and within family body odor signatures, as different types of skin bacteria and their concentration affect the formation of the latter (40). It has been established that the human microbiome can be transmitted indirectly (via substrates such as paper, cotton or glass) and directly via physical contact (41), implying that if family members spend a lot of time together, their microbiomes become more similar. Thus, the effect of family-specific patterns in component overlaps may be due to some parents being (physically) closer with their children. However, current research lacks quantification of how physical contact like cuddling affects body odor similarity. In addition, the formation of body odor may be modulated by hormonal and physiological processes, including stress-related changes in endocrine activity (42). In our sample, we did neither control for cuddling habits nor for current stress levels of the odor donors. Also dietary aspects impact body odor (19,43), which we standardized by handling dietary restrictions to the donors. Nevertheless, individuals from the same family and household, like the donor families from our study, might overlap even more in diet (e.g. sharing meals regularly), potentially accounting for part of component overlap and family body odor similarity. Additionally, offspring may share dietary preferences more strongly with one parent than the other, resulting in partial alignment of odor profiles within specific parent-offspring dyads, while odor differences to the remaining family members may reflect divergent dietary habits.

In the present study, we took two axillary TD samples from each participant for the chemical analysis. Although instrumental variability in GC/MS is typically low (26), intra- and inter-individual variability can be substantial. Hence, Weiß et al. (44) emphasized the importance of taking at least two to three samples per individual. Due to feasibility when assessing human subjects, we decided on the lower threshold of two replicates. Hence, future studies should invest into higher sample sizes and multiple samples per individual.

Both our perceptual and chemical studies provided evidence that perceptual and chemical similarity of body odor relates to kinship. With these parallel lines of results, we expected to also find a correlation between both domains. Accordingly, we observed a small but positive significant relationship of perceptual and chemical similarity. In general, measuring associations between chemical signatures and their percept is quite complex (45–47), as perception is highly individual (e.g. with respect to ethnicity, personality, see (48,49) and often leads to only small effects (46). It further presents several methodological constraints that were also pertinent in our study: sampling methods for perception - and chemical studies were different, i.e. cotton pads vs. TD tubes, which are currently the state-of-the-art methods (28,50). Both methods overlap in capturing a proportion of volatile compounds, but differ in recapture-capacity of semi-and non-volatile compounds (25). Moreover, cotton pad sampling integrated body odor over a 12-h collection period, whereas TD tube sampling captured a momentary snapshot of body odor at sampling time. Finally, TD tube sampling was performed twice, while cotton pad sampling occurred only once. Yet, despite these methodological limitations, we observed a significant link, indicating a promising direction for future research. More refined methods may help characterize the chemical underpinnings of perceived body odor similarity in humans more precisely. The statistical approach of employing a machine learning based predictor represents a fruitful strategy to maximize information yield without losing relevant information in prior aggregating, or dimension reduction, especially regarding the high dimensionality of chemical data. This is paralleled by recent approaches, targeting the intersection of chemistry and human perception by machine-learning (51,52).

Overall, further methodological research is needed to explore adequate matching of perceptual and chemical methods. In the present study, we mostly sampled volatile and semi-volatile compounds with the TD tube approach, while non-volatile compounds may also contribute significantly to the olfactory perception of volatiles present (53). Hence, it could be beneficial to extend chemical sampling in future studies. For example, Owsienko et al. (54) used direct contact sampling with pretreated cotton pads, followed by solvent extraction and GC-O/GC-MS when sampling body odor.

Such methodological advances may enable a more comprehensive characterization of those chemical features that contribute to perceived body odor similarity. Establishing this link between chemical composition and perception is an important step toward understanding how molecular characteristics of body odor translate into perceptual, behavioral, and neurobiological responses. In the longer term, these insights may provide candidate targets for clinical interventions strengthening family relationships.

## Conclusion

To our knowledge this study is the first to jointly analyze perceptual and chemical markers of olfactory kin recognition in biological family structures of humans. Sex and relatedness predicted perceived as well as chemical odor similarity. Integrative analyses revealed a small but significant link between perception and chemistry. While this first integrative analysis of body odor profiles of biological families extends earlier human studies on the matter, future research is needed to increase our understanding of the relationship between chemical signatures and percept in human body odor.

## Acknowledgements

This project was funded by the Free State of Saxony and the German Federal Ministry of Education and Research (BMBF; TUD FOSTER A3-2022 to KH). The chemical analysis was partially funded by the European Fund for Regional Structure Development, EFRE (“Europe funds Saxony”), grant no. 100195810 to AW as well as by the Leipzig University, with technical assistance from the MS-UL MS Core Facility at the Institute of Analytical Chemistry, Faculty of Chemistry, Leipzig University. This work was also funded by the Deutsche Forschungsgemeinschaft (DFG, German Research Foundation) – GRK 2950 – Project-ID 509922606 (BH).

## References

1. Hamilton WD. The genetical evolution of social behaviour. I. Journal of Theoretical Biology. 1. Juli 1964;7(1):1– 16. doi:10.1016/0022-5193(64)90038-4

2. Chapais B, Savard L, Gauthier C. Kin selection and the distribution of altruism in relation to degree of kinship in Japanese macaques (Macaca fuscata). Behav Ecol Sociobiol. 1. Mai 2001;49(6):6. doi:10.1007/s002650100335

3. Park JH, Schaller M, Van Vugt M. Psychology of Human Kin Recognition: Heuristic Cues, Erroneous Inferences, and Their Implications. Review of General Psychology. 1. September 2008;12(3):3. doi:10.1037/1089-2680.12.3.215

4. Widdig A, Nürnberg P, Krawczak M, Streich WJ, Bercovitch FB. Paternal relatedness and age proximity regulate social relationships among adult female rhesus macaques. Proc Natl Acad Sci U S A. 20. November 2001;98(24):13769– 73. doi:10.1073/pnas.241210198 PubMed PMID: 11698652; PubMed Central PMCID: PMC61116.

5. Lehmann L, Perrin N. Inbreeding avoidance through kin recognition: choosy females boost male dispersal. Am Nat. November 2003;162(5):638–52. doi:10.1086/378823 PubMed PMID: 14618541.

6. Widdig A. Paternal kin discrimination: the evidence and likely mechanisms. Biological Reviews. 2007;82(2):319–34. doi:10.1111/j.1469-185X.2007.00011.x

7. DeBruine LM, Smith FG, Jones BC, Craig Roberts S, Petrie M, Spector TD. Kin recognition signals in adult faces. Vision Research. 1. Januar 2009;49(1):38–43. doi:10.1016/j.visres.2008.09.025

8. Gustafsson E, Levréro F, Reby D, Mathevon N. Fathers are just as good as mothers at recognizing the cries of their baby. Nat Commun. 2013;4:1698. doi:10.1038/ncomms2713 PubMed PMID: 23591865.

9. Henkel S, Setchell JM. Group and kin recognition via olfactory cues in chimpanzees (Pan troglodytes). Proceedings of the Royal Society B: Biological Sciences. 24. Oktober 2018;285(1889):20181527. doi:10.1098/rspb.2018.1527

10. Schäfer L, Croy I. An integrative review: Human chemosensory communication in the parent-child relationship. Neurosci Biobehav Rev. Oktober 2023;153:105336. doi:10.1016/j.neubiorev.2023.105336 PubMed PMID: 37527693.

11. Weiß BM, Kücklich M, Thomsen R, Henkel S, Jänig S, Kulik L, u. a. Chemical composition of axillary odorants reflects social and individual attributes in rhesus macaques. Behav Ecol Sociobiol. 28. März 2018;72(4):65. doi:10.1007/s00265-018-2479-5

12. Charpentier M, Boulet M, Drea C. Smelling right: The scent of male lemurs advertises genetic quality and relatedness. Molecular ecology. 1. Juli 2008;17:3225–33. doi:10.1111/j.1365-294X.2008.03831.x

13. Porter RH, Cernoch JM, Balogh RD. Odor signatures and kin recognition. Physiology & Behavior. 1. März 1985;34(3):445–8. doi:10.1016/0031-9384(85)90210-0

14. Kuhn F, Natsch A. Body odour of monozygotic human twins: a common pattern of odorant carboxylic acids released by a bacterial aminoacylase from axilla secretions contributing to an inherited body odour type. Journal of The Royal Society Interface. 6. April 2009;6(33):377–92. doi:10.1098/rsif.2008.0223

15. Penn DJ, Oberzaucher E, Grammer K, Fischer G, Soini HA, Wiesler D, u. a. Individual and gender fingerprints in human body odour. J R Soc Interface. 22. April 2007;4(13):13. doi:10.1098/rsif.2006.0182

16. Schäfer L, Sorokowska A, Sauter J, Schmidt AH, Croy I. Body odours as a chemosignal in the mother–child relationship: new insights based on an human leucocyte antigen-genotyped family cohort. Philosophical Transactions of the Royal Society B: Biological Sciences. 8. Juni 2020;375(1800):1800. doi:10.1098/rstb.2019.0266

17. Watzlawik M. Die Erfassung des Pubertätsstatus anhand der Pubertal Development Scale: Erste Schritte zur Evaluation einer deutschen Übersetzung: Diagnostica: Vol 55, No 1. Diagnostica. 2009;55(1):55–65. doi:10.1026/0012-1924.55.1.55

18. Havlicek J, Fialová J, Roberts S. Individual Variation in Body Odor. In. 2017. S. 125–6. doi:10.1007/978-3-319-26932-0_50

19. Havlicek J, Lenochova P. The Effect of Meat Consumption on Body Odor Attractiveness. Chem Senses. 1. Oktober 2006;31(8):8. doi:10.1093/chemse/bjl017

20. Croy I, Mohr T, Weidner K, Hummel T, Junge-Hoffmeister J. Mother-child bonding is associated with the maternal perception of the child’s body odor. Physiology & Behavior. 1. Januar 2019;198:151–7. doi:10.1016/j.physbeh.2018.09.014

21. Hierl K, Croy I, Schäfer L. Body Odours Sampled at Different Body Sites in Infants and Mothers—A Comparison of Olfactory Perception. Brain Sciences. Juni 2021;11(6):6. doi:10.3390/brainsci11060820

22. Lenochova P, Roberts SC, Havlicek J. Methods of Human Body Odor Sampling: The Effect of Freezing. Chem Senses. 1. Februar 2009;34(2):2. doi:10.1093/chemse/bjn067

23. Lötsch J, Ultsch A, Hummel T. How Many and Which Odor Identification Items Are Needed to Establish Normal Olfactory Function? Chem Senses. Mai 2016;41(4):4. doi:10.1093/chemse/bjw006 PubMed PMID: 26857742.

24. Python [Internet]. Python Software Foundation; 2025. Verfügbar unter: https://www.python.org/

25. Kücklich M, Möller M, Marcillo A, Einspanier A, Weiß BM, Birkemeyer C, u. a. Different methods for volatile sampling in mammals. PLOS ONE. 25. August 2017;12(8):e0183440. doi:10.1371/journal.pone.0183440

26. Marcillo A, Jakimovska V, Widdig A, Birkemeyer C. Comparison of two common adsorption materials for thermal desorption gas chromatography – mass spectrometry of biogenic volatile organic compounds. Journal of Chromatography A. 8. September 2017;1514:16–28. doi:10.1016/j.chroma.2017.07.005

27. Stein SE. An integrated method for spectrum extraction and compound identification from gas chromatography/mass spectrometry data. Journal of the American Society for Mass Spectrometry. 1. August 1999;10(8):770–81. doi:10.1016/S1044-0305(99)00047-1

28. Kücklich M, Weiß BM, Birkemeyer C, Einspanier A, Widdig A. Chemical cues of female fertility states in a non-human primate. Sci Rep. 23. September 2019;9(1):1. doi:10.1038/s41598-019-50063-w

29. R Core Team [Internet]. Vienna; 2021. (R Foundation for Statistical Computing). Verfügbar unter: https://www.R-project.org

30. Oksanen J, Simpson G, Blanchet FG, Kindt R, Legendre P, Minchin P, u. a. vegan community ecology package version 2.6-2 April 2022. 2022.

31. Ricotta C, Podani J. On some properties of the Bray-Curtis dissimilarity and their ecological meaning. Ecological Complexity. 1. September 2017;31:201– 5. doi:10.1016/j.ecocom.2017.07.003

32. Chung H, Beretvas SN. The impact of ignoring multiple membership data structures in multilevel models. Br J Math Stat Psychol. Mai 2012;65(2):185– 200. doi:10.1111/j.2044-8317.2011.02023.x PubMed PMID: 21732931.

33. Bürkner PC. brms: An R Package for Bayesian Multilevel Models using Stan. Journal of Statistical Software. 2017;(80(1)):1–28. doi:10.18637/jss.v080.i01

34. Taleb F, Vasco M, Ribeiro AH, Björkman M, Kragic D. Can Transformers Smell Like Humans?

35. Tom G, Tian Ser C, Rajaonson EM, Lo S, Suk Park H, Lee BK, u. a. Does this smell the same? Learning representations of olfactory mixtures using inductive biases. Mach Learn: Sci Technol. 30. September 2025;6(3):035063. doi:10.1088/2632-2153/adfffc

36. Goodfellow I, Bengio Y, Courville A. Deep Learning [Internet]. MIT Press; 2016. Verfügbar unter: http://www.deeplearningbook.org

37. Balbona JV, Kim Y, Keller MC. The estimation of environmental and genetic parental influences. Development and Psychopathology. Dezember 2022;34(5):1876–86. doi:10.1017/S0954579422000761

38. Visscher PM, Medland SE, Ferreira MAR, Morley KI, Zhu G, Cornes BK, u. a. Assumption-Free Estimation of Heritability from Genome-Wide Identity-by-Descent Sharing between Full Siblings. PLOS Genetics. 24. März 2006;2(3):e41. doi:10.1371/journal.pgen.0020041

39. Boulet M, Charpentier MJ, Drea CM. Decoding an olfactory mechanism of kin recognition and inbreeding avoidance in a primate. BMC Evolutionary Biology. 3. Dezember 2009;9(1):281. doi:10.1186/1471-2148-9-281

40. Troccaz M, Gaïa N, Beccucci S, Schrenzel J, Cayeux I, Starkenmann C, u. a. Mapping axillary microbiota responsible for body odours using a culture-independent approach. Microbiome. 24. Januar 2015;3(1):1. doi:10.1186/s40168-014-0064-3

41. Neckovic A, van Oorschot RAH, Szkuta B, Durdle A. Investigation of direct and indirect transfer of microbiomes between individuals. Forensic Science International: Gen tics. 1. Ärz 2020;45:102212. doi:10.1016/j.fsigen.2019.102212

42. Katsuyama M, Narita T, Nakashima M, Kusaba K, Ochiai M, Kunizawa N, u. a. How emotional changes affect skin odor and its impact on others. PLOS ONE. 30. Juni 2022;17(6):e0270457. doi:10.1371/journal.pone.0270457

43. Hepper PG. The discrimination of human odour by the dog. Perception. 1988;17(4):549–54. doi:10.1068/p170549 PubMed PMID: 3244526.

44. Weiß BM, Marcillo A, Manser M, Holland R, Birkemeyer C, Widdig A. A non-invasive method for sampling the body odour of mammals. Methods in Ecology and Evolution. 2018;9(2):420–9. doi:10.1111/2041-210X.12888

45. Keller A, Gerkin RC, Guan Y, Dhurandhar A, Turu G, Szalai B, u. a. Predicting human olfactory perception from chemical features of odor molecules. Science. 24. Februar 2017;355(6327):820–6. doi:10.1126/science.aal2014

46. Kowalewski J, Huynh B, Ray A. A System-Wide Understanding of the Human Olfactory Percept Chemical Space. Chem Senses. 1. Januar 2021;46:bjab007. doi:10.1093/chemse/bjab007

47. Li H, Panwar B, Omenn GS, Guan Y. Accurate prediction of personalized olfactory perception from large-scale chemoinformatic features. Gigascience. 1. Februar 2018;7(2):gix127. doi:10.1093/gigascience/gix127

48. Parma V, Redolfi N, Alho L, Rocha M, Ferreira J, Silva CF, u. a. Ethnic influences on the perceptual properties of human chemosignals. Physiology & Behavior. 15. Oktober 2019;210:112544. doi:10.1016/j.physbeh.2019.05.005

49. Sorokowska A, Groyecka A, Karwowski M, Frackowiak T, Lansford JE, Ahmadi K, u. a. Global Study of Social Odor Awareness. Chemical Senses. 24. August 2018;43(7):503–13. doi:10.1093/chemse/bjy038

50. Zetzsche M, Weiß BM, Kücklich M, Stern J, Birkemeyer C, Widdig A, u. a. Combined perceptual and chemical analyses show no compelling evidence for ovulatory cycle shifts in women’s axillary odour. Proceedings of the Royal Society B: Biological Sciences. 24. Juli 2024;291(2027):20232712. doi:10.1098/rspb.2023.2712

51. Mersha M, Lam K, Wood J, AlShami AK, Kalita J. Explainable artificial intelligence: survey of needs, techniques, direction. September applications, and future Neurocomputing. 28. 2024;599:128111. doi:10.1016/j.neucom.2024.128111

52. Schreurs M, Piampongsant S, Roncoroni M, Cool L, Herrera-Malaver B, Vanderaa C, u. a. Predicting and improving complex beer flavor through machine learning. Nat Commun. 26. März 2024;15(1):2368. doi:10.1038/s41467-024-46346-0

53. Tomasino E, Song M, Fuentes C. Odor Perception Interactions between Free Monoterpene Isomers and Wine Composition of Pinot Gris Wines. J Agric Food Chem. 11. März 2020;68(10):3220–7. doi:10.1021/acs.jafc.9b07505

54. Owsienko D, Goppelt L, Hierl K, Schäfer L, Croy I, Loos HM. Body odor samples from infants and post-pubertal children differ in their volatile profiles. Commun Chem. 21. März 2024;7:53. doi:10.1038/s42004-024-01131-4 PubMed PMID: 38514840; PubMed Central PMCID: PMC10957943.

