## Supplement for "The chemistry of families: Linking body odor composition to olfactory perception"

**Title: Supplementary methods**

Caption: Supplementary information regarding all methodological aspects of the study that is reported in main manuscript can be found here.

**1.** **GENERAL METHODS**

a) BO sampling procedure

At an initial appointment, BO donor families were equipped with a study kit containing all materials required to collect BO samples. This included two cotton pads, each stored in an airtight wide-mouth pharmacy jar, medical adhesive tape, fragrance-free body soap (https://www.eubos.de/hautpflege/basis-pflege/fluessig-wasch-dusch-nachfuellbeutel), fragrance-free washing powder (https://www.dm.de/denkmit-vollwaschmittel-pulver-ultra-sensitive-p4066447328820.html), a disposable razor, a written declaration on the study kit, behavioral requirements, and a declaration on the General Data Protection Regulation. Participants received study instructions and provided written informed consent.

Participants were asked to adhere to the following protocol 48 hours before sampling with cotton pads and 48 hours before the TD-Tube samplings at T1 and T2. Strongly seasoned food, onions, garlic, leek, asparagus, cabbage, and alcohol should be avoided. In addition, participants were asked to follow a vegetarian or vegan diet (1,2). Places with a strong odor, such as swimming pools, were to be avoided, and no perfumed products were to be used. Instead, we provided participants with perfume-free body soap for showering, and perfume-free laundry detergent for their bedlinen, towels, and clothing that was worn prior to and during BO sampling/ experimental sessions in our laboratory.

*BO sampling with cotton pads for sensory analysis.* In order to obtain BO samples with cotton pads (stimuli for intra-familiar and third-party BO rating) participants were asked to adhere to a standardized behavioral protocol that has been proven in numerous studies (3–5). They were instructed to shower with the fragrance-free body soap and, if desired, to shave their armpits. They were asked about the hair status of their armpits at sampling time later. Cotton pads were attached with medical adhesive tape in the evening and worn under both armpits for 12 hours throughout the night. Only clothing that had previously been washed with the perfume-free laundry detergent was to be worn. In addition, bed linen in which the participants slept during sampling were to be washed with the same perfume-free detergent. At the end of sampling period, cotton pads were placed back in the airtight pharmacy jars. On the same day, BO samples were frozen at -25°C at the University Hospital Dresden and stored until they were needed for sensory experiments. Lenochova and colleagues (6) demonstrated that freezing BO samples does not affect their quality.

*TD-Tube sampling for chemical analyses.* At the same appointment collecting the cotton BO pads back in the lab, BO components were additionally sampled using Thermal desorption (TD) tubes (stainless steel TD tubes, Supelco 1/4 in. × 3 1/2 in., Supelco, Bellefont, USA) (T1). A second sampling point was conducted after at least three days (*M* = 19.60, *SD* = 18.62, min = 3, max = 61 days). These adsorbent traps contained two polymers (.1 g Tenax TA and .2 g XAD-2, Sigma Aldrich) to capture the volatile (VOC) and semi-volatile organic compounds (sVOC). Paring of these two polymers ensures comprehensive coverage and sensitivity for sampling VOC (substances that evaporate easily, pass into the air and can be directly detected by the olfactory sense) and sVOC (substances that are less volatile than VOCs and are more likely to be found directly on surfaces such as skin or fur; (7). For sample collection, TD tubes were connected to an air pump (BiVOC2, Holbach) with a plastic hose. Each TD tube was positioned as close as possible to the axillary of the participant without touching their skin. The air pump was set to 1.5L/min flow rate and produced a constant air flow until .5 L of air was collected. Sampling took place under laboratory conditions to reduce environmental influences. For each participant, order of left and right axilla sampling tool place in a randomized order on T1/T2 respectively. Experimenter wore lab coat, mask and nitrile gloves to minimize contamination. 15 families, each consisting of four family members, were sampled on both axilla sides, resulting in *N*=120 TD tubes. Additionally, 11 room blanks and 3 analytical blanks were collected as control of contaminations. The room blanks only contained air of the sampling room and the analytical blanks were transported and stored like the other tubes but without using them. This served controlling for contamination caused by transport, storage and chemical compounds in the room.

Before samples were collected, each TD tube was cleaned under a constant stream of nitrogen in a thermal conditioner (TD Clean-Cube, Scientific Instruments Manufacturer) for 120 minutes and was closed with Swagelok brass caps on both sides immediately afterwards. Furthermore, the cleaned tubes were wrapped in aluminum foil, stored in airtight plastic bags before and after sampling, to avoid contamination.

**2.** **PERCEPTION STUDY**

Olfactory ability of all human raters was tested on the basis of a psychophysiological test using a screening version of the Sniffin' Sticks. Participants were asked to identify three odors (cinnamon, banana, fish smell) in a 4-alternative forced-choice task. This test is sufficient to assess normosmia and ensures reliable results with a sensitivity of 80.4% and a specificity of 84.3% (8). If no correct identification of the three odors was achieved, the complete identification test consisting of 16 odors was performed (exclusion criterion < 12 correctly identified odors). In addition, current odor impairments (e.g. rhinitis) were queried at the beginning of the test in order to postpone the test in the event of an impairment.

**Assessment by third parties**

Rating procedure:

Third party raters were invited to evaluate perceived similarity of BO samples. Participants were informed about the aim and purpose of the study and asked for the written consent. They were asked to refrain from wearing perfume and other strong-smelling care products, and smoking on the day of the experiment, and from eating and drinking coffee 1 hour before the experiment. BO samples were presented according to the same standardized procedure as described earlier (cotton pad BO samples were used, as well). After presenting two anchor BOs, participants were asked to rate the similarity of two BO samples ("How similar are these two BO samples?") on a visual analogue scale from 0 to 100 (0 = not at all, 100 = very). Related and unrelated dyad-combinations from a total of 3 families (*n* = 12 individuals) matched for sex of the children were presented to each participant. Possible dyad combinations were sibling-dyads, unrelated adolescent dyads, parent-child dyads, and unrelated adult-adolescent dyads. The order of these combinations, and the order of the first dyad sample to be presented were randomized. A total of 20 pairs were to be assessed. In a second step, participants were asked to match an initially presented BO sample with one of three other samples whose donor was related to the donor of the initial sample (3 alternative forced choice paradigm; "Which of these three BOs belongs to the initially presented BO?"). In total, participants were asked to identify brothers, sisters, mother- daughter-, mother-son-, father-daughter- and father-son-dyads (olfactory phenotype matching) in 12 phenotype-matching trials.

**3. CHEMICAL STUDY**

Chemical analyses

GC-MS: Given the number of samples and blanks, evaluation was split into two batches. At the beginning and end of both batches, three GC-MS blanks were run to clear out possible residues from previous measurements. Additionally, one GC-MS blank was run after every tenth sample, and the room blanks as well as the analytical blanks were analyzed randomly among test samples in both batches. Due to methodological problems, 6 samples from 6 different subjects were lost, resulting in a sample size of *N* = 114.

Chemical profiling: Chemical profiles were processed by a semi-automated method. AMDIS (v. 2.65, 9) was initially used to obtain a peak list for each sample for automated signal deconvolution and peak picking. Subsequently, the individual peak lists were merged and all recurring peaks with similar retention times (RTs) were grouped into RT ranges. These ranges underwent manual correction by considering the consistency of their RTs and specific m/z ratios. RT ranges needed to occur in at least 4 out of all 114 samples to be considered for further analysis. Retention time and corresponding substance-specific m/z has been assigned for 269 retention time ranges. Afterwards, resulting compound library was applied to the entire dataset using Shimadzu GCMS Browser software, searching for each library entry based on RTs and most distinctive m/z ratios.

Finally, obtained data were cleaned to minimize the influence of contaminations and analytical procedures by excluding peak areas outside the retention time range (± 50%) and peak areas below 1000. Additionally, constantly appearing compounds in analytical blanks were excluded if the peak area was equal or higher than in the human samples, and compounds occurring more than three times higher in room blanks than in samples were excluded as well. As a result, the final dataset was reduced to 130 substances.

Statistical analyses

2) In depth assessment of chemical similarity: Further questions on the similarities of BO profiles were investigated using multiple membership multilevel models (MMMM), based on pairwise Bray-Curtis (dis)similarities. These models were also based on pairwise Bray-Curtis (dis)similarities calculated using the package ‘vegan’, version 2.6-2 (10) from standardized, log(x+1)-transformed peak areas which were computed previously. The continuous predictors age and sampling date were z-transformed to a mean of 0 and an SD of 1 before model fitting. Given that similarity scores follow beta distribution, all models were calculated using a Bayesian approach in the package ´brms´ (version 2.21.0, 11), resulting estimates refer to the posterior distribution. The ‘brms’ package implements MMMM with beta distribution using the function ‘vif’ from the R package ‘car’ (version 3.1-2, 12) to assess collinearity between the predictor variables. Rhat values and the effective sample sizes (ESS) were examined to ensure model convergence. The Rhat values confirm the degree of variance within and between the chains. The ESS values serve to assess both the number of independent samples within the central area of the distribution (Bulk_ESS) and the extent to which the extreme areas of the distribution are covered (Tail_ESS). These two values should ideally be above 1000 to ensure good convergence.

First, we examined effects of relatedness on BC-similarity in i) parent-child versus unrelated dyads, and ii) siblings versus unrelated dyads. Second, effects of sex on BC-similarity was examined in iii) same-sex parent-child versus opposite-sex parent-child dyads, and iv) sister-dyads versus brother-dyads.

i) Parent-child vs. control-dyads (relatedness)

A subset consisting of all parent-offspring dyads and all unrelated adult-child dyads in the data set was created to test the impact of relatedness of BO profiles. When formulating this model, the warning message of high divergent transition after warmup indicated that the model was not correctly exploring the distribution, resulting in distorted estimates (Stan Development Team, 2022-03-10). Therefore, all participants with only one sample needed to be excluded for this analysis (resulting in a samples size of *N*=108 in this analysis). In MMMM, dyadic Bray-Curtis similarities were fitted as continuous beta-distributed response variable and the test predictor for this model was family ID. This dataset contained same sex and opposite sex dyads. Sex (same)was included as fixed effect control predictor as well as age, sampling date and GC-MS batch. GCMS file name (unique number per sample) and sample ID (respective person to the sample) were fitted as multi-membership terms as they have levels that are present in more than one variable. As there were two samples of most participants, a unique ID was created for each combination of participants before, to ensure correct integration in the model. The GC-MS file name, sample ID and unique ID were included in the model as random effects. For the GC-MS-sample number and sample ID, random slopes were needed for family, sex, age, sampling date and batch because of sufficient variations. Random slopes were necessary for the unique ID within sampling date and GC-MS batch, however, family, sex and age were not variable in the context of unique ID and therefore there were no random slopes for these parameters. The model converged with all Rhat values deviating from 1 by <.01 and with ESS>1000 at 12000 post-warmup draws.

ii) Siblings vs. control-dyads (relatedness)

The model fitting for the similarity comparison between same-sex siblings and unrelated same-sex children was identical to the first model (i). However, for this analysis the subset contained only same sex sibling-dyads and unrelated same-sex control-dyads. In this model, also participants with only one sample were included. In contrast to (i), there was no random slope for GC-MS batch in the unique ID, as the subsample size was smaller and thus the batch variation was insufficient. This model converged with all Rhat values deviating from 1 by <.01 and with ESS>1000 at 8000 post-warmup draws. There was a warning of 1 divergent transition after warmup but Rhat und ESS values were sufficiently high to move forward (Stan Development Team, 2022-03-10).

iii) Same-sex vs. opposite-sex parent-child dyads (sex)

Similarities of same sex parent-offspring dyads and opposite sex parent-offspring dyads were calculated to investigate the effect of sex. This model included the participants with only one sample as well. Sex (same) was fitted here as a test predictor, as the model analyzed the impact of sex on the body odor similarities between parent-offspring dyads. Age, sampling date and GC-MS batch were fitted as fixed effects control predictor and the unique ID, GC-MS file name and sample ID as random effects as in the models above. In this model was no need for random slopes because of the limited subsample size. This model converged with all Rhat values deviating from 1 by <.01 and with ESS > 765 (most ESS > 1000) at 4000 post-warmup draws.

iv) Male vs. female sibling-dyads (sex)

In this analysis male vs. female sibling dyads were compared and the model was fitted similar to (iii) with only one modification: As this subset only included sibling dyads, the sample ID and unique ID were identical and consequently, sample ID was omitted in this MMMM. Furthermore, in each family were either two sisters or two brothers who were compared with other siblings of same sex. Consequently, the data set contains only same-sex dyads, with sex specified as female or male. Sex (female) was integrated as test predictor. This model converged with all Rhat values deviating from 1 by <.01 and with ESS > 948 (most ESS > 1000) at 4000 post-warmup draws.

**4. References**

1.	Havlicek J, Fialová J, Roberts S. Individual Variation in Body Odor. In 2017. S. 125–6.

2.	Havlicek J, Lenochova P. The Effect of Meat Consumption on Body Odor Attractiveness. Chem Senses. 1. Oktober 2006;31(8):747–52.

3.	Croy I, Mohr T, Weidner K, Hummel T, Junge-Hoffmeister J. Mother-child bonding is associated with the maternal perception of the child’s body odor. Physiology & Behavior. 1. Januar 2019;198:151–7.

4.	Hierl K, Croy I, Schäfer L. Body Odours Sampled at Different Body Sites in Infants and Mothers—A Comparison of Olfactory Perception. Brain Sciences. Juni 2021;11(6):820.

5.	Schäfer L, Sorokowska A, Sauter J, Schmidt AH, Croy I. Body odours as a chemosignal in the mother–child relationship: new insights based on an human leucocyte antigen-genotyped family cohort. Philosophical Transactions of the Royal Society B: Biological Sciences. 8. Juni 2020;375(1800):20190266.

6.	Lenochova P, Roberts SC, Havlicek J. Methods of Human Body Odor Sampling: The Effect of Freezing. Chem Senses. 1. Februar 2009;34(2):127–38.

7.	Kücklich M, Möller M, Marcillo A, Einspanier A, Weiß BM, Birkemeyer C, u. a. Different methods for volatile sampling in mammals. PLOS ONE. 25. August 2017;12(8):e0183440.

8.	Lötsch J, Ultsch A, Hummel T. How Many and Which Odor Identification Items Are Needed to Establish Normal Olfactory Function? Chem Senses. Mai 2016;41(4):339–44.

9.	Stein SE. An integrated method for spectrum extraction and compound identification from gas chromatography/mass spectrometry data. Journal of the American Society for Mass Spectrometry. 1. August 1999;10(8):770–81.

10.	Oksanen J, Simpson G, Blanchet FG, Kindt R, Legendre P, Minchin P, u. a. vegan community ecology package version 2.6-2 April 2022. 2022.

11.	Bürkner PC. brms: An R Package for Bayesian Multilevel Models using Stan. Journal of Statistical Software. 2017;(80(1)):1–28.

12.	Fox J, Weisberg S. An R Companion to Applied Regression. Sage Publications. 2018;(1).
